# Characterization of a Unique Spontaneous Calcifying Cell Line (CJ): a Novel Tool for the Study of Ectopic Calcification

**DOI:** 10.1101/2024.01.05.574307

**Authors:** Nobutaka Ida, Yoshihisa Yamane

## Abstract

**Purpose:** Due to the lack of an appropriate in vitro evaluation system, there is no effective prevention or treatment for ectopic calcification diseases (ECD). We obtained canine renal adenocarcinoma cells (CJ cells) that spontaneously form large amounts of calcified precipitates (CaP-ppt) and examined whether they could serve as a model for the early stages of ectopic calcification (EC).

**Methods:** Dispersed cells obtained by collagenase-DNase digestion were cultured in 10% FBS, antibiotic-containing DMEM-high glucose medium (standard medium). CaP-ppt was stained with Alizarin Red (AR) and colorimetrically quantified after acid extraction. Cell volume was determined by Crystal violet (CV) staining followed by extraction and colorimetric quantification. Calcium (Ca) and phosphate (PO_4_) were determined with a commercial kit to obtain the Ca/P ratio. Screening of anti-CaP-ppt compounds was performed in the same 96-well plate in the order of cell culture ⇒ CaP-ppt quantification ⇒ cell quantification and evaluated by IC_50_ value.

**Results:** CJ cells produced large amounts of CaP-ppt on standard medium alone without external phosphate addition; CaP-ppt formation was not accompanied by cell death, but on the contrary, CaP-ppt increased at physiological pH values around pH 7.4 due to active cell metabolism. The Ca^2+^ and PO_4_^3-^ partitioning kinetics to CaP-ppt were found, and the Ca/P ratio of CaP-ppt was stable at 1.35. The anti-Cap-ppt effects of bisphosphonates and all-trans retinoic acid (ATRA) were also confirmed in this CJ cell CaP-ppt system.

**Discussion:** There was doubt about the vascular calcifying cell model with the addition of high concentrations of phosphoric acid, but there were no suitable alternative cells. Spontaneously calcifying CJ cells provide a fundamental solution to this problem. Anti-CaP-ppt screening also eliminates the need for medium exchange, thus saving labor and cost. The Ca/P ratio of CaP-ppt in CJ cells is 1.35, the same as that of amorphous Ca phosphate (ACP), corresponding to the early (reversible) stage of EC. Therefore, it has favorable conditions as an evaluation system for drug discovery.

**Conclusion:** CJ cells, which calcify at physiological phosphate concentrations (0.9 mM) in standard media, are useful and novel research material for basic and preventive studies of ECD and for drug development studies.

## 1 : Introduction

Ectopic calcification (EC) is a phenomenon that results in mineral deposition in soft tissues that do not normally calcify, as opposed to physiological calcification in bones, cartilage, and teeth. In recent years, the phenomenon of EC has been widely reported in atherosclerosis, aortic valve calcification, urinary tract stones, age-related macular degeneration, breast cancer, prostate cancer, and Alzheimer’s disease, for which there are no effective treatments or therapeutic agents, and calcification inhibition is attracting attention as a new therapeutic target for these intractable diseases[1–3]。

In particular, "cardiovascular calcification (CVC)" and "renal stone disease (RSD)" are the most common ectopic calcification diseases with many patients, and there are many similarities between them (Table 1). Furthermore, these are multifactorial diseases that are complexly influenced by diverse pathophysiological, genetic, and environmental factors, and despite the vast amount of basic and clinical research on both, there are no effective preventive or therapeutic agents for CVC [4–6] and RSD [7–9]. One of the reasons for this is the lack of appropriate materials for mechanistic elucidation and an efficient evaluation system to screen candidate compounds for therapeutic agents. Moreover, recently, doubts have emerged regarding the conventional cellular-assessment method of EC itself, using vesicular smooth muscle cells (VSMC).

**Table 1 :** Elements of cardiovascular calcification and urinary lithiasis as ectopic calcification.

| Item | Cardiovascular calcification | Kidney stone |
| --- | --- | --- |
| Calcification type | Calcium Phosphate | Calcium oxalate, Calcium Phosphate |
| Risk factors | Aging, Obesity, Diabetes, Chronic renal disease, Metabolic syndrome | Aging, Obesity, Diabetes, Hypertension, Metabolic syndrome |
| Effective therapeutic agents | None | None |
| Effective physical treatment | Yes (Rotablator, IVUS<sup>a)</sup> etc.) | Yes (TUL<sup>b)</sup>, ESWL<sup>c)</sup> etc.) |
| Calcification site | vascular endometrium (intima, inflammatory) and/or vascular middle layer (medial, non-inflammatory) | Intraurethral and/or Interstitium (non-inflammatory) |
| Luminal structure | Yes | Yes |
| Fluids in the lumen | Blood | Urine |
| Plaque formation | Yes (Atheroma plaque) | Yes (Randall's plaque) |
| Fluid on Plaque | Blood flow (intravascular lumen) | Urine flow (renal pelvis) |
| Exfoliation plaque-induced symptoms | Myocardial infarction, Cerebral infarction | Ureteral obstruction |
| Plaque components | ACP<sup>d)</sup>, HAP<sup>e)</sup>, Collagen, Laminin, Cells | ACP, HAP, Collagen, Cells |

a) IVUS: Intravascular ultrasound, <sup>b)</sup>TUL: Transurethral ureter lithotomy, <sup>c)</sup>ESWL: Extracorporeal shock wave, <sup>d)</sup>ACP: amorphous calcium phosphate, <sup>e)</sup>HAP: hydroxyapatite

The study of model cells for calcification began in the field of bone research. Assuming that "the phenomenon of calcification itself is common," whether physiological or pathological, it is natural to try to learn from prior bone biology. Bone cell lines are especially useful due to their unlimited availability and high experimental reproducibility. For example, in the mouse osteoblast MC3T3-E1 strain [10, 11], high concentrations (5-10 mM) of *β*-glycerophosphate (β-GP) with ascorbic acid produce large amounts of calcified deposits, which has become the standard method for inducing calcification in other osteoblast cell lines [12]. In regenerative medicine, the addition of 10 mM *β*-GP to bone induction from tissue stem cells (e.g. bone marrow stromal cells and adipose stem cells) has been used in an in vitro 2D culture system [13] and a 3D-printed hydroxyapatite (HAP) scaffold perfusion culture system [14].

Aging, diabetes, and chronic kidney disease (CKD) are risk factors for CVC, and elevated blood phosphate levels in patients with advanced stage CKD and end-stage renal disease (ESRD) are well known[15, 16]. Therefore, it was a natural progression that the effect of *β*-GP addition obtained in bone studies would be applied to CVC studies. Indeed, the addition of 2 mM phosphate, twice the physiological concentration (1 mM), induced a large amount of calcium phosphate precipitation (hereafter referred to as CaP-ppt) and increased expression of the osteogenic genes Runx2 and osteocalcin in a human VSMC culture system [17]. The impact of this increase in bone marker gene expression was so great that the theory of transformation of VSMC into osteoblast-like cells and the addition of *β*-GP or inorganic phosphate quickly became popular in CVC studies *in vitro* and is now the standard method for inducing calcification[18–20].

On the other hand, in the field of bone metabolism [21–23] and bone formation in regenerative medicine [24, 25], there were continuing reports of doubt that the CaP-ppt produced in high concentrations of *β*-GP cannot be said to be from normal bone tissue.

In 2015, Hortells *et al*. challenged the unnaturalness of this highly phosphate-added experimental model using rat VSMC in a very precise and important experiment [26].The results showed that CaP-ppt formation in the presence of high concentrations of phosphate (2 and 3 mM) was a physicochemical precipitation caused by a shift in culture medium pH toward alkaline and was completely unrelated to the actual vascular calcification that occurs in the body. On the other hand, the authors suggested that the local pH in the body (vascular tunica media smooth muscle) must be alkalized for this to be a model for arterial calcification[26, 27]. However, no reports of local alkalinization have been found so far.

Furthermore, Hortells *et al*. and Millan *et al*. calculated that even when severe hyperphosphatemia (2.5 mM) is reached, phosphate concentration does not reach saturation (S(ACP)=0.53) with amorphous calcium phosphate (ACP) at acidotic blood pH values in CKD patients. In other words, the basic premise that the Ca x P product value based on hyperphosphatemia values in CKD is the cause of vascular calcification, S(ACP)>1.0, the theoretical value of supersaturation, is not valid [26, 27]. This is consistent with O’Neill’s assertion that EC cannot be explained by the "Ca x P product theory" [28]. The above *in vitro* experimental results and *in vivo* thermodynamic analysis strongly suggest that EC due to CKD and ESRD may not be caused by hyperphosphatemia.

In 2021, Front. Cell Dev. Biol. asked 28 leading experts in their fields of research to contribute to a review of the mechanisms, risk factors, pathogenesis, and prevention of ECD. The journal sought contributions from 28 leading experts in their fields of research on the mechanisms, risk factors, pathogenesis, and prevention of ECD. The journal received excellent original papers and review articles, but none of the articles provided updates on available or promising treatment strategies [1].

Calcification based on CKD hyperphosphatemia in advanced stages is exceptional within the group of ECD [26]。In fact, hyperphosphatemia has not been observed in other groups of ectopic calcifying diseases such as urinary tract stones, age-related macular degeneration, breast cancer, prostate cancer, and Alzheimer’s disease. Thus, we should now turn our attention to ECD occurring at normal phosphate concentrations" [29] and the corresponding *in vitro* experimental models.

Spontaneous calcification under normal phosphate concentration has already been reported in two strains of bovine aortic SMC (CVC strain) [30, 31] and canine distal tubular cells (MDCK strain) [32, 33]. However, they have a major weakness in that the calcification signal is extremely weak and have not been utilized as a model for vascular calcification or renal calculi. In this study, we show the unique properties of canine renal adenocarcinoma-derived cells (CJ cells) that form spontaneous and massive calcified precipitates under physiological conditions that overcome this weakness. To establish the canine and feline tumor panel, we have already obtained about 40 passage-ready cell lines from over 200 surgical specimens (the protocols have been submitted to and approved by the Ethics Committee of the Animal Clinical Research Foundation). And the CJ cell line was obtained by fortunate chance in the process.

## 2 : Materials and Methods

### 2 – 1 : Materials

MDCK cells were kindly provided by Dr. Jobu Ito (College of Medicine, Tokai University, Japan). 4.5 g/L glucose-containing DMEM (DMEM-HG), Alizarin red S, Crystal violet All-trans retinoic acid, and Colcemid were purchased from FUJIFULM-Wako Pure Chemical, Osaka Japan. Collagenase (type I) and Deoxyribonuclease I (DNase I) were purchased from Sigma, NY, USA. Fetal Bovine Serum (FBS) was purchased from Biowest, Nuaillé, France.

Penicillin G and streptomycin were obtained from Meiji Seika, Tokyo, Japan. Etidronate and Alendronate were obtained from Tokyo Kasei Kogyo, Tokyo, Japan. Cell culture flasks and manufacturers used were as follows: for 25 cm^2^ flasks, model no. 430639 Corning NY USA, CELLSTAR® 690-170 Greiner Bio-One GmbH Frickenhausen Germany, model no. 3103-025; for 75 cm^2^ flasks, model no. 3123-075 IWAKI AGC Technoglass Corporation (Shizuoka, Japan) or model no. MS-23250 SUMILON® Sumitomo Bakelite Corporation (Tokyo, Japan) were used. All multi-well plates (model numbers omitted) were SUMILON® Sumitomo Bakelite Co Ltd. Tokyo Japan.

### 2 – 2 : Preparation of CJ cells

Cells were prepared from adenocarcinoma tissue of the left kidney surgically removed from a 3-year-old female Chihuahua dog. The tumor tissue was cut into small sections, transferred to a mixture of 0.1% (w/v) collagenase Type I and 0.01% (v/v) DNase I dissolved in 40 ml HBSS(+) and stirred at 37℃ for 60 minutes. After filtering through a 100 μm stainless steel mesh, the cells were washed by centrifugation at 1200 rpm, 4℃, for 5 min and dispersed in new PBS(-) again for the second cell washing by the centrifugation. The cells were dispersed in DMEM-HG containing 10% FBS, 100 u/ml penicillin, 100 μg/ml streptomycin, 4.5 g/L glucose (hereafter referred to as standard medium or SM). Approximately 5 × 10^5^ cells/ml SM, seeded into uncoated Corning 25 cm^2^ flasks, and cultured in an incubator at 37℃, 5% CO_2_ and 95% air. The cells were designated "CJ".

### 2 – 3 : Cell passage and chromosome counting

Before reaching confluency, CJ cells were passaged. After medium removal and PBS(-) washing, CJ cells were detached from the flasks by addition of 0.25% trypsin-1mM EDTA-4Na (Fujifilm Wako Pure Chemicals) and diluted with fresh SM at a sprit ratio 1 : 4 or 1 : 8 and transferred into fresh flasks. Chromosome analysis of CJ cells was performed according to conventional methods [15].

### 2 – 4 : Morphometrics, pH measurement, viable cell evaluation and cell fixation

An Olympus CKX-41 equipped with a centering phase difference slider (IX2-SL) with ring slit (IX2-SLPH2) was used for microscopic observation. In each test, pH measurement of the culture supernatant was performed before viable cell determination and cell fixation, if necessary. After photographing the color tone of the medium in the 96-well plate, the medium was immediately dropped onto the flat electrode of a pH meter, LAQUAtwin® pH33B (HORIBA Scientific, Kyoto, Japan), and the pH was measured. In this test, 150 μl was dropped from the supernatant culture medium (about 200 μl) in the well onto the electrode (flat sensor section), and each sample was measured within 1 minute. Viable cells were evaluated using the trypan blue dye exclusion method with a blood cell counting board or CCK-8 (Cell Counting Kit-8, Dojindo). The CCK-8 assay was performed according to the vendor’s instructions. Unless otherwise specified, the cells were washed twice with PBS(-), fixed with 95% ethyl alcohol (EtOH, 100 μl/well) for 30 minutes at room temperature, and stored until various staining. Details of the various experimental conditions are specified in the Results section of each study or in the figure descriptions.

### 2 – 5 : Detection of calcified deposits

Ca in CaP-ppt was detected by alizarin red S (AR) staining. EtOH-fixed samples were stained with 1% (w/v) AR stain for 30 min at room temperature, washed with water to remove excess dye, dried at room temperature, and photographed from the back of the culture vessel. If necessary, cells were counterstained with 1% light green after AR staining. von Kossa staining was performed by fixing cells in 95% EtOH or 100% methyl alcohol (MeOH), washing once with distilled water (dH_2_O), and adding 5% (w/v) silver nitrate (Fujifilm Wako Pure Chemicals). The culture dish was irradiated with tabletop fluorescent light for 30 minutes, washed again with dH_2_O, and then 5% sodium thiosulfate was added and allowed to react for 3 minutes. After the sodium thiosulfate was removed, the culture dishes were dried and visualized after three dH_2_O washes. Photographs were taken at each step if necessary.

### 2 – 6 : Determination of calcium and cells by dye staining methods

Unless otherwise specified, staining quantitative tests for CaP-ppt and test cells were performed as follows: CJ cells were sown in 96-well plates at 2 x 10^4^ cells/200 μl SM /well. On the day of determination, the cells were fixed described above and stained with 60 μl/well of 1% (w/v) alizarin red S for 30 min at room temperature. AR-stained CaP-ppt was dissolved in 5% formic acid (150 µl/well), and Ca content was measured as absorbance at OD 450 nm. The same plates were subsequently used to estimate the cell volume (organic matter) by CV staining and extraction. That is, after removing the above AR extract solution, the plates were rinsed thoroughly with water and once dried. Then, 60 μl/well of 0.05 % (w/v) crystal violet (CV) solution was added, stained for 30 min at room temperature, and excess CV dye was removed by washing with water. CV-stained cells were treated with 150 μ/well of the extracting mixture (ethylene glycol: 70% EtOH: citric acid = 50 : 50 : 1) and the absorbance value at 590 nm was measured.

### 2 – 7 : Alkaline phosphatase (ALPase) staining

Detection of ALPase in CJ and MDCK cells was performed using the azo dye coupling method (Burstone 1958) with slight modifications. After medium removal, cells were washed twice with 5 mM Hepes buffered saline and fixed in cold MeOH containing 10% (v/v) formaldehyde and 0.01% (v/v) glacial acetic acid for 10 seconds. Substrate solutions were prepared by dissolving 50 mg naphthol AS-BI phosphate (Sigma-Aldrich) and 50 mg Fast Blue RT (Sigma-Aldrich) in 100 ml of 50 mM 2-amino-2-methyl-1,3-propanediol buffer (pH 9.8). Enzyme reactions were performed by adding 1 ml/well of substrate solution and incubating at room temperature for 30 min.

### 2 – 8 : Quantification of calcium (Ca) and phosphorus (P)

CJ cells were seeded in 96-well plates at 2 x 10^4^ cells/200 μl SM/well, and samples were collected, fixed in the wells, and treated with drying every 2 days. At the time of treatment, wells other than the treated wells were first covered with film sheets to prevent evaporation and contamination, and the culture supernatant was collected from the treated wells, sealed, and stored aseptically at room temperature. The treated wells were then washed twice with 5 mM Hepes buffered saline (200 μl/well), fixed with 95% EtOH (100 μl/well) for 30 minutes at room temperature, the fixing solution was removed, and after air-drying, the film sheets were removed, and the plates were returned to the CO_2_ incubator for continued culture. This sampling procedure was repeated every 2 days until day 14. After all sampling was completed, the sample wells of the air-dried plates were injected with 200 μl/well of 0.6N HCl, sealed with a film sheet, and left at room temperature for 24 hours to elute calcium and phosphate. Calcium was determined with the Calcium E-test Wako assay kit (Wako Pure Chemical, Osaka, Japan) based on the MXB method, and phosphate with the BIOMOL®

GREEN assay kit (Enzo Life Science, Inc. NY USA). Both absorbance measurements were performed at OD 590 nm using Benchmark (Bio-Rad).

### 2 – 9 : Validation of CJ cell HTS system

CJ cells diluted to 10^5^ cells/ml with SM were sown at 200 μl/well in all wells of 96-well plates except rows 1 and 12, and culture was started. Validation tests were performed two times (first and second trials).

### 2 – 10 : Screening by IC_50_ values for anti-calcification and cytotoxicity of test compounds

The test compounds were evaluated using a combination of the AR-stained extraction method (calcification effect) and CV-stained extraction method (cytotoxic effect). The test compounds were dissolved in 100% DMSO and stored at -20℃ as stock solutions unless otherwise noted. On day-0, CJ cells dispersed in "screening medium" were seeded in all wells of 96 well plates (2 × 10^4^ cells/200 μl/well) except for the first row. On day-2, the test substance was added. That is, 200 μl of the highest concentration of the compound solution diluted in screening medium was directly added to the culture wells of CJ cells in the second row (400 μl/well) and mixed, then 200 μl of the mixture was transferred to the next row and mixed, and then a 2-fold step dilution was performed until the 11th row, The culture was resumed. Judgment was made on day-8, and the % values of AR and CV staining of the test substance relative to Cell control (100%) and Blank (0%) were calculated. IC_50_ values for anti-CaP effect and cytotoxicity were calculated using GraphPad Prism (Version 7, Graphpad Software, Inc. La Jolla, CA, USA).

### 2 – 11 : Statistical analysis

All measurement results are expressed as mean ± standard deviation. GraphPad Prism 7 (La Jolla, CA, USA) was used for all statistical analyses. Tests for two groups with no correspondence were performed by *t*-test. One-way ANOVA was used for analyses between multiple groups. Two-way-ANOVA was used to analyze multiple treatments at multiple time points. Dunnett’s multiple comparison test was used for paired control comparisons, and Tukey’s multiple comparison test was used for comparisons between all groups. Sidak’s multiple comparisons were also performed for comparisons between time points in the two groups; *p* < 0.05 was considered significantly different.

## 3 : Results

### 3 – 1 : CJ cells form large amounts of deposits spontaneously

During passaging of CJ cells, we noticed a large amount of sediment at the bottom of the "backup flasks" left in the CO_2_ incubator (Fig. 1a-2). No precipitate formation or turbidity of the culture medium was observed in the SM-only flasks used to check for contamination (Fig. 1a-1). CJ cells grew well in SM, forming colonies composed of epithelial-like cells (Fig. 1b), with a mean doubling time of 23.2 hr in log-growth phase(Fig. 1c). The chromosome number of CJ cells was 2n = 78 ± 16 (n = 11), approximating the haploid number of 78 for normal canine cells (Fig. 1f). Immediately after confluent arrival, CJ cells showed distinct intercellular borders (Fig. 1c), and dome formation also began around confluence (Fig. 3c). Subsequently, CaP-ppt formation began with abrupt dome retraction, and by the second week after passage, a white tone "island" composed of CaP-ppt and a "sea" portion composed only of cells were formed (Fig. 1a-2, 1d). The CaP-ppt portion consisted of brownish-white particles of various sizes and their aggregates, and iridescent (pinkish in the photograph) smaller-sized microspheres of about 1∼2 μm in diameter were also observed (Fig. 1d). CaP-ppt stained favorably to alizarin red S stain and von Kossa stain, and cell parts to Giemsa stain and crystal violet stain (Fig. 1g). After a longer period, CaP-ppt exhibited various morphologies, including aggregation, fusion, and enlargement, based on spherical particles of various sizes (Fig. 2). The mean size of the spherical particles, measured from observations at 100x with a phase contrast microscope, was 5.0 ± 2.2 μm (Mean ± SD). In passages 142 PDLs, CJ cells were confirmed by a third-party institution to be mycoplasma free.

**Figure 1:**
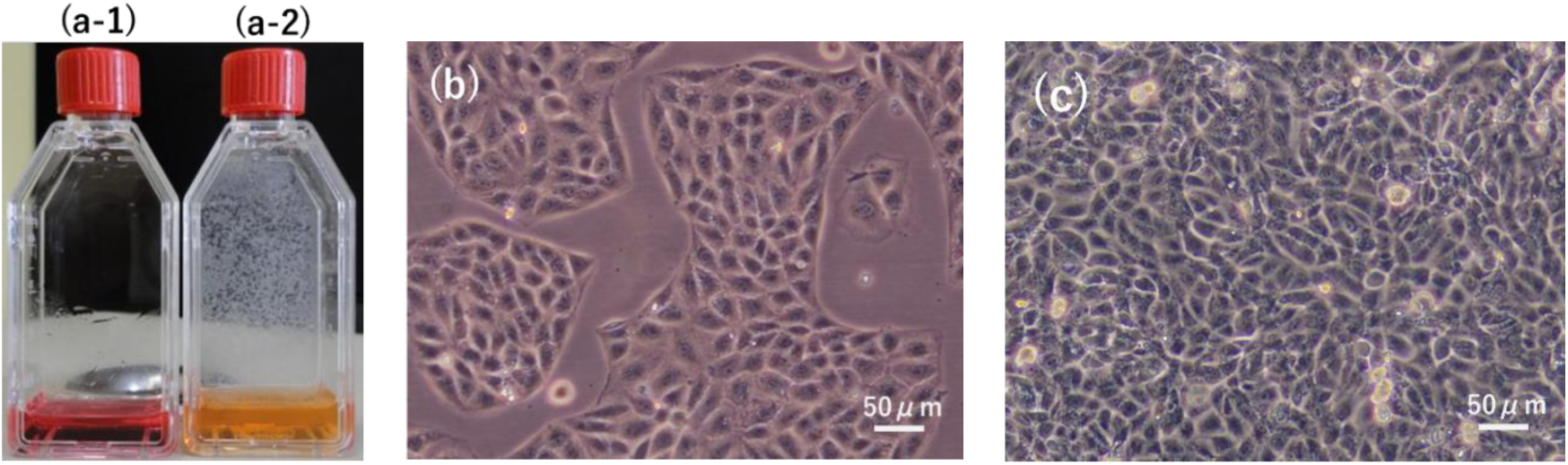
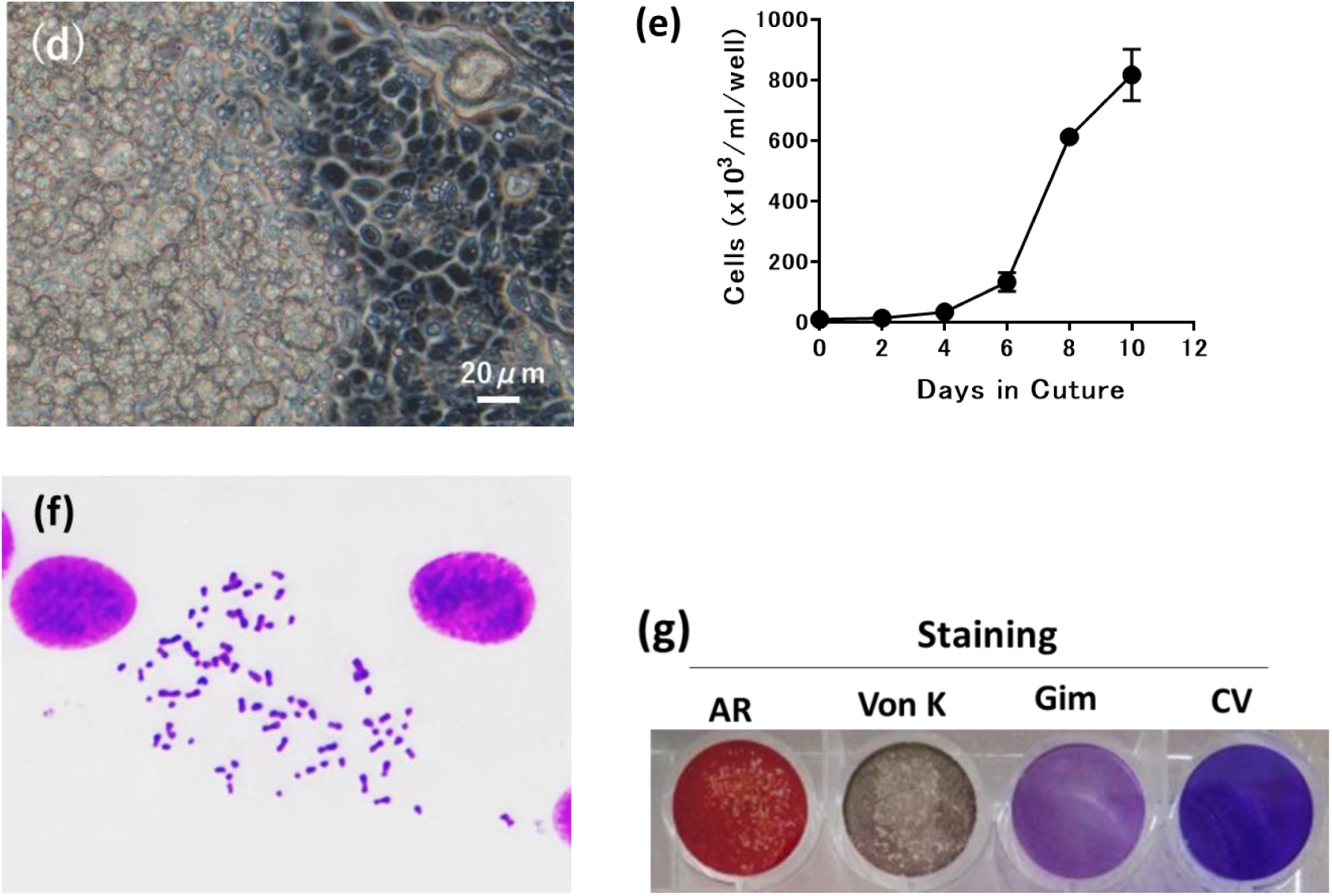
CJ cells show strong. (**a-1**) SM alone, (**a-2**) CaP-ppt of CJ cells. (**b**) Colonies of logarithmically growing CJ cells. (**c**) CJ cell sheet immediately after reaching Confluent. No CaP-ppt formation is seen yet. (**d**) CaP particles of various sizes produced by CJ cells after 2 months of culture with weekly medium changes. (**e**) CJ cell growth curve in 24-well plates, sown at 1x 10^4^ cells/ml/well on day-0, cells were detached with 0.25% Trypsin-1mM EDTA every 2 days, and viable cell counts were calculated on a blood cell counting board (each point Mean±SD, n=4). (**f**) Chromosome image of CJ cells. (**g**) Histochemical staining of CJ cells after CaP-ppt formation. On day-0, 2 x 10^5^ cells/ml of CJ cells were sown in 24-well plates at 1 m/well, and on day 10 of culture, four stains were performed in individual wells: AR: alizarin red S stain, von K: von Kossa stain, Gim: Giemsa stain, CV: crystal violet stain.

**Figure 2:**
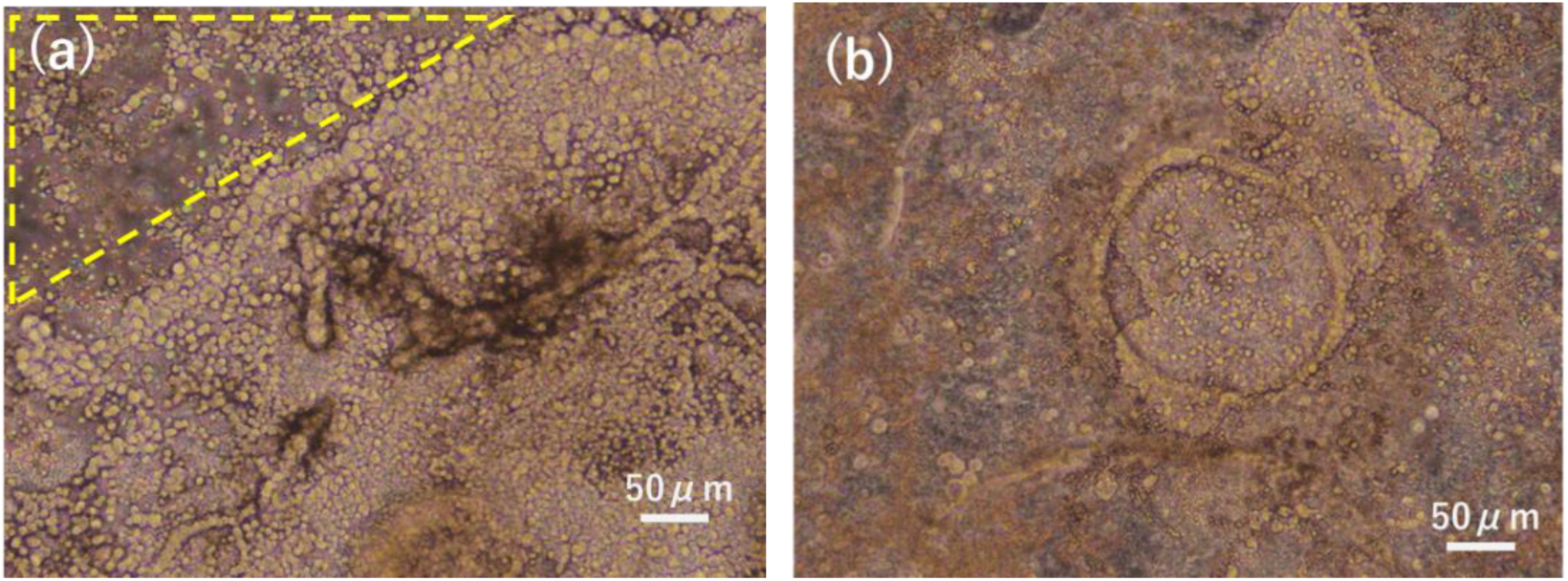
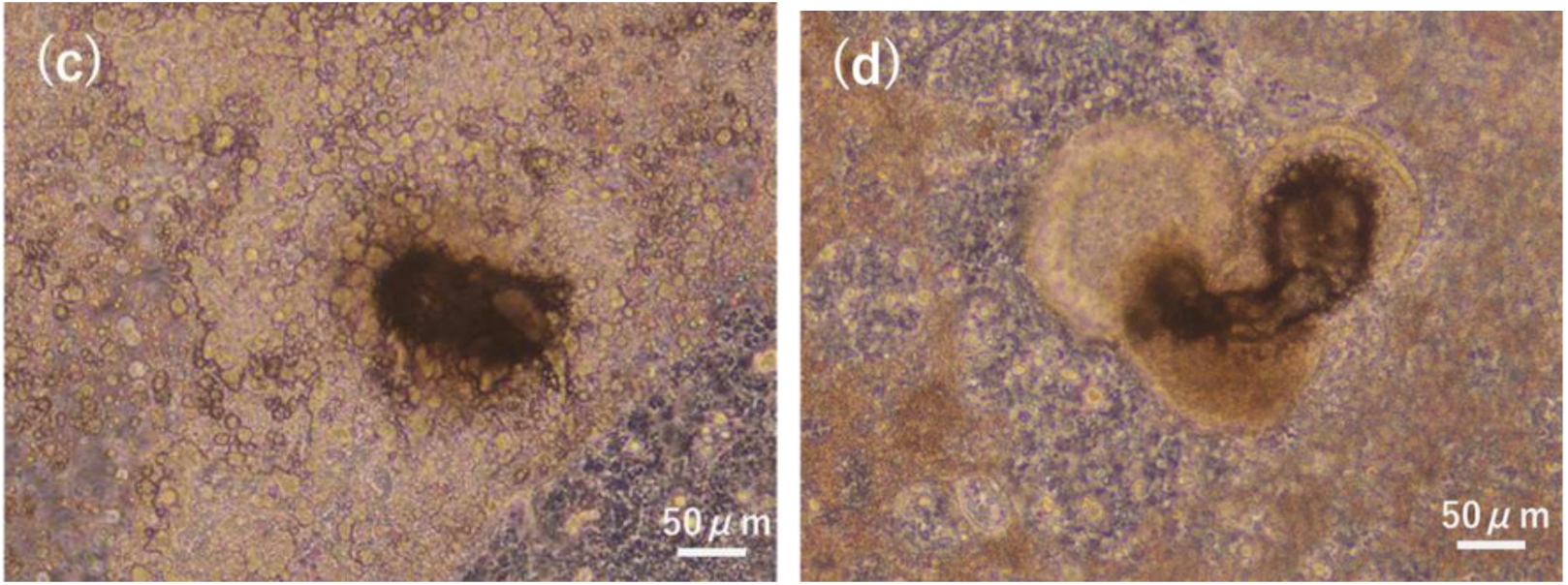
Various shapes of CaP-ppt. formed in Corning 25 cm^2^ flasks after 2 months of weekly medium changes. (**a**) Mutual aggregation and linear to branched root-like, (**b**) Plate-like growth or circular linear caldera or crater structure by microparticles, (**c**) Growth as aggregates in the center of plate-like growth, and (**d**) Small CaP particles and their aggregation inside spheroid-like structure. Particle size was measured in the area circled by the yellow dotted triangle in the upper left of photo (a). That is, the photo (a) was electronically enlarged and printed in B5 size, the outer circumference of the spherical particles was transferred onto tracing paper, and the diameter was measured with a caliper and converted from the projected scale size. The diameter of a total of 90 particles (Min. 2.3 μm to Max. 14.3 μm) measured was 5.0 ± 2.2 μm (Mean ± SD).

**Figure 3:**
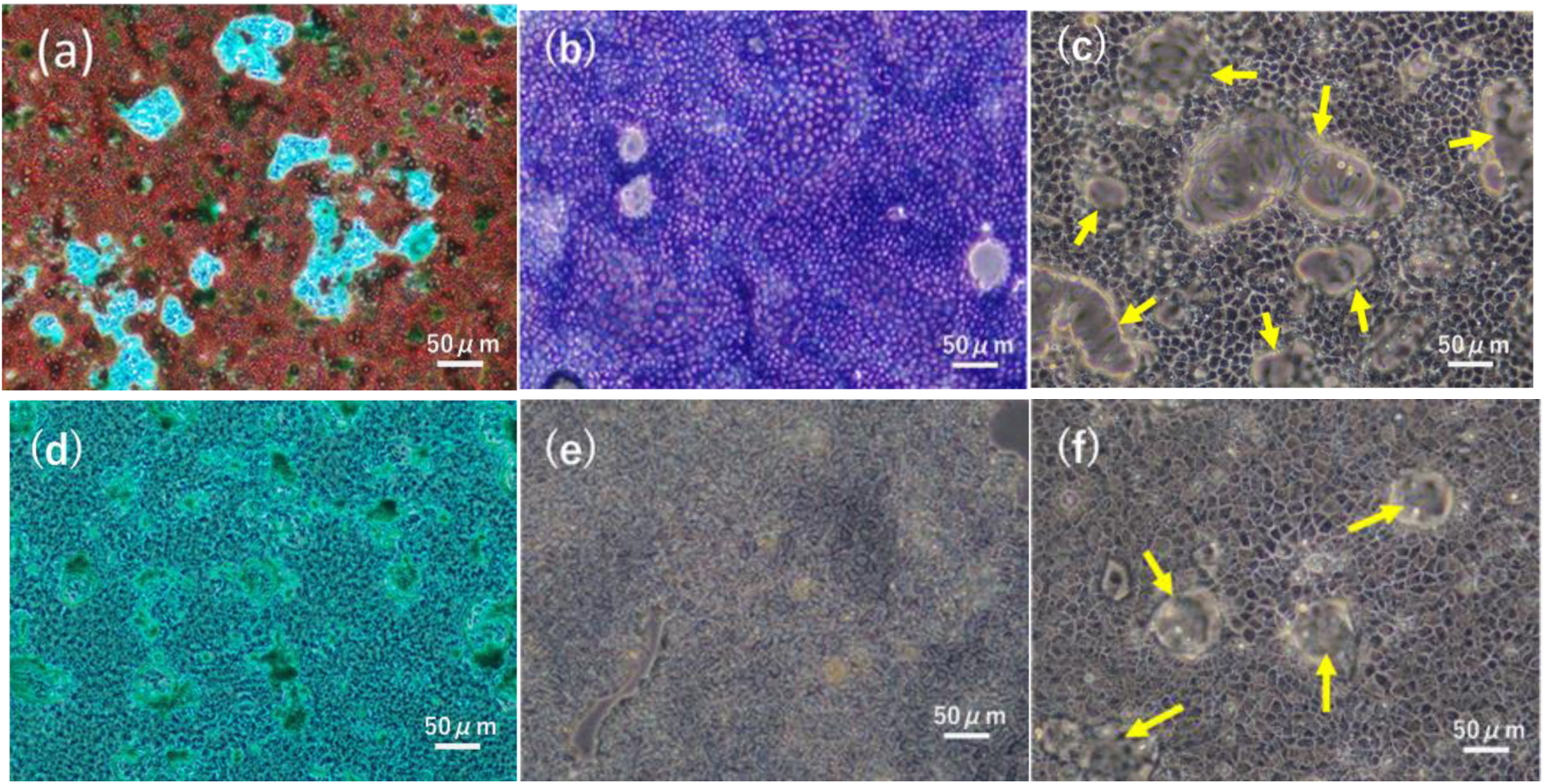
CJ cells characterize glandular cells derived primarily from the proximal tubules of the kidney. (**a**) through (**c**) are CJ cells and (**d**) through (**f**) are MDCK cells. (a) and (d) are double staining of cells with AR staining for Ca-ppt detection (red stained area) and light green (LG) staining (green stained area), (b) and (e) are ALPase staining, (c) and (f) are photographs of dome formation (yellow arrows). Test cells were diluted to 1 x 10^5^cells/ml with SM and sown in 6-well plates at 3ml/well to start culture (day-0). Double staining with AR/LG dye (a, d) was performed on day 8 and ALPase staining on day 5 (b, e). The remaining two wells for dome observation had the medium removed on day-6 and were photographed in PBS(-).

### 3 – 2 : The ability of CJ cells to form CaP-ppt is an order of magnitude greater than that of known MDCK cells

Spontaneous calcification was reported in MDCK cells derived from the distal tubules of canine kidney [32, 33] and compared to CJ cells in terms of calcification ability; CJ cells were stained in a pattern of "sea island structures" consisting of abundant calcified deposits stained red by AR dye and green stained cells only by LG dye (Fig. 3a). In contrast, MDCK cells showed only green-stained cells and no red-stained calcified areas (Fig. 3b). After reaching confluency, both cells showed dome formation characteristic of glandular cells (Fig. 3c, f), and ALPase staining showed high activity in CJ cells (Fig. 3c), whereas it was very low in MDCK cells (Fig. 3e). Based on the presence of high ALPase activity, a marker of proximal tubular cells [34, 35], the CJ cells obtained in this study were presumed to be mainly of proximal tubular origin.

### 3 – 3 : FBS is required for calcification of CJ cells

The effect of FBS on calcification was compared between CJ and NDCK cells. The amount of CaP-ppt formation in CJ cells increased with the concentration of FBS added, reaching a plateau at FBS concentrations of 6% to 10%; CaP-ppt formation in MDCK cells was not observed at different FBS concentrations (Fig. 4a). On the other hand, cell proliferation showed sufficient cell proliferation effects at concentrations of 1% to 2% FBS for both CJ and MDCK cells. Interestingly, cell number comparison by CV staining [36] showed that CJ cells stained significantly stronger than MDCK cells (Fig. 4b).

**Figure 4:**
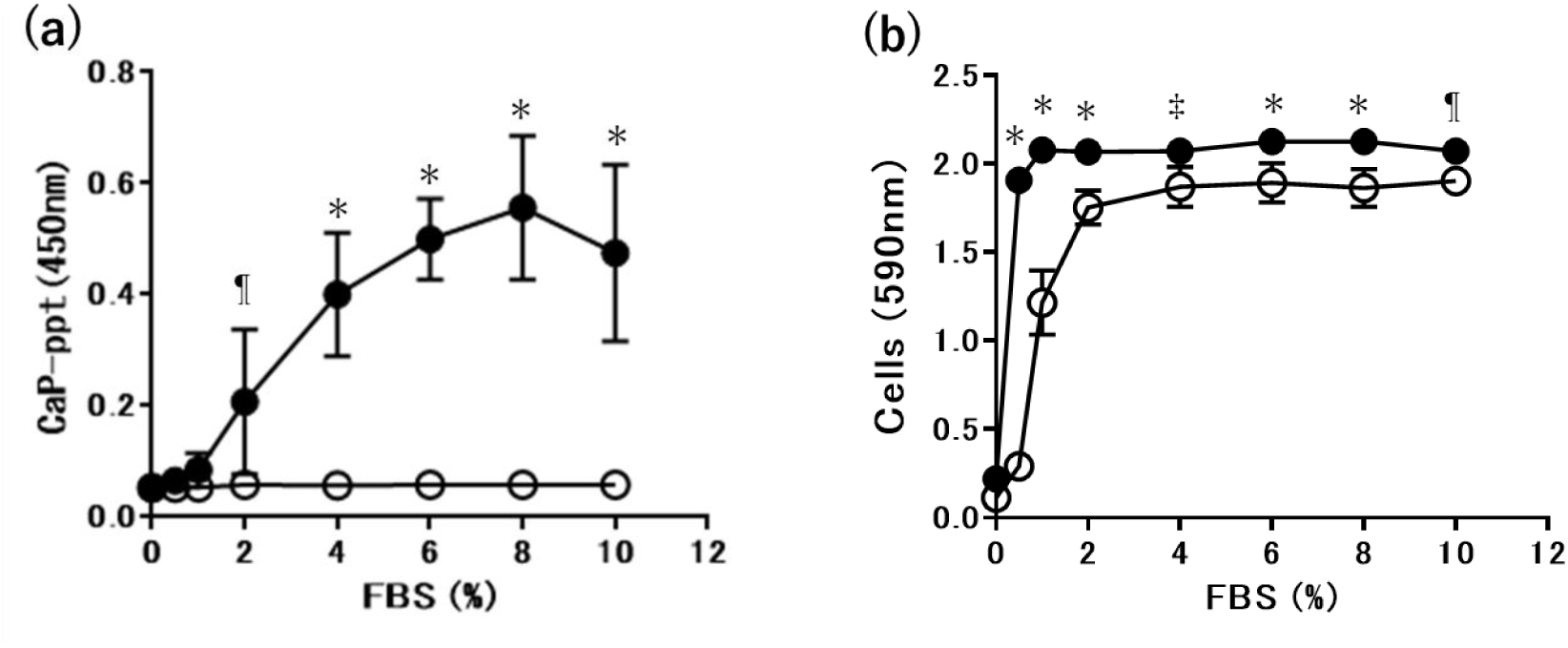
FBS requirement for Cap-ppt formation in CJ and MDCK cells. CJ cells (●) and MDCK cells (○) were detached by trypsin-EDTA treatment, thoroughly dispersed in SM in 15 ml centrifuge tubes, and centrifuged to obtain cell sediment after counting viable cells. To this cell sediment, 10 ml/tube of PBS(-) was added and stirred, and PBS centrifugal washing was performed twice. To the washed cells, DMEM-HG medium containing the indicated final concentration of FBS, prepared separately, was added, and well dispersed to prepare a test cell solution of 1 × 10^5^ cells/ml. On day-0, cultures were started in 96-well plates at 10^4^ cells/100 μl/well. AR staining and CV staining were performed consecutively on the same plate on day-8, and OD values were determined as described in the Methods section. (a) Amount of CaP-ppt by AR staining and (b) Amount of cells by CV staining. Statistical analysis was performed by 2-way ANOVA with Sidak’s multiple-comparison test for each FBS concentration of the corresponding MDCK cells. ¶*p*<0.005, ‡*p*<0.0005, \**p*<0.0001.

### 3 – 4 : CaP-ppt formation kinetics and material balance in CJ cells

There has been no long-term follow-up of the transfer of supernatant Ca and phosphate ions to CaP precipitates in the formation of CaP-ppt in cell culture systems from the viewpoint of material balance. The reason for this is that in other culture systems, the medium was changed every 2 to 3 days. On the other hand, CJ cells can be cultured for a long period (more than 20 days) without changing the culture medium. Therefore, we measured the transfer and distribution of initial amounts of Ca and phosphorus (P) to the culture medium and precipitates in a closed culture system with CJ cells.

As expected, the amount of Ca in the culture medium decreased with time, and in parallel, the amount of Ca in the CaP-ppt formed at the bottom of the culture vessel increased (Fig. 5a). Similarly, the amount of phosphate in the culture medium of CJ cells decreased with time, while the amount of phosphate in the precipitates increased (Fig. 5b), confirming that the main body of CaP-ppt in CJ cells is Ca phosphate by this quantification method in addition to AR staining. The intersections of the descending and ascending curves of Ca and P concentrations in the supernatant and precipitate in this experiment were 6-7 days after the start of culture. The total amount of Ca and inorganic phosphorus partitioned into the supernatant and precipitate was within the theoretical range after day-6. However, on day-2 and day-4, both were significantly lower than the theoretical values.

**Figure 5:**
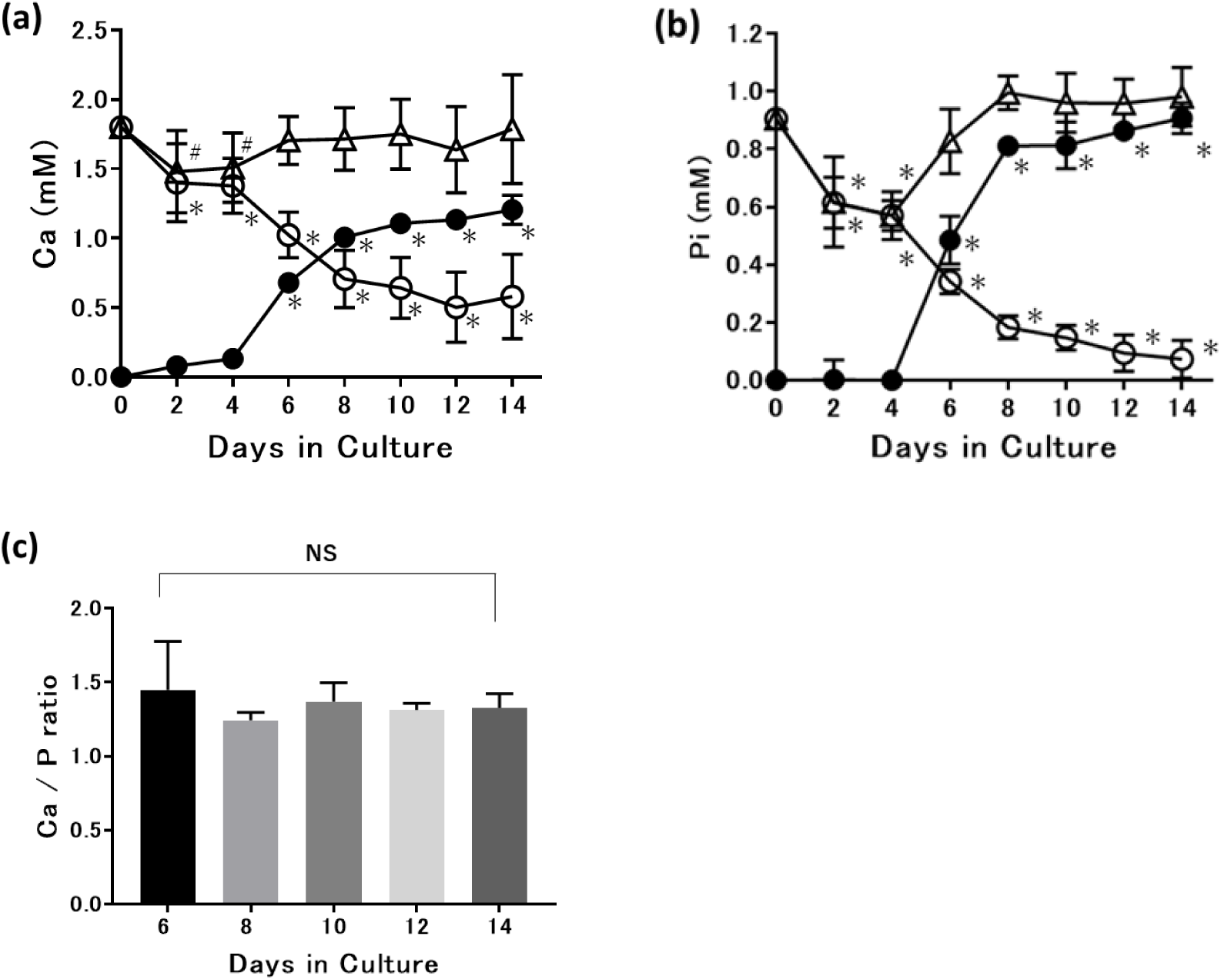
Calcification of CJ cells is due to Ca and phosphorus in the culture medium. Experiments were performed as described in the Methods section. No medium was changed during the experiment. Each point in the figure is shown as Mean±SD (n=8). (a) Ca^2+^ partitioning into supernatant (○) and precipitate (●) in CJ cell cultures. (b) PO4^3-^ partitioning into supernatant (○) and precipitate (●) in CJ cell cultures. (△) is the sum of supernatant and precipitate. Statistical analysis was performed by 1-way ANOVA with Dunnett’s multiple-comparison test. #*p*<0.05 vs day-0, \**p*<0.0001 vs day-0. (c) Ca/P ratio calculated from CaP-ppt. Statistical analysis was performed by 1-way ANOVA with Tukey’s multiple-comparison test; NS=not significant; values around 1.35 indicated by amorphous calcium phosphate persisted from day 6 to day 14.

The Ca/P ratio of CaP-ppt was 1.34 ± 0.12 (Mean ± SD), corresponding to the amorphous Ca phosphate [29] value between days 6 and 14, consistent with the theoretical value for material balance, and was not significantly different between time points (Fig. 5c).

### 3 – 5 : Active calcification of CJ cells proceeding in the physiological pH range

All experiments were performed in 96-well plates, and the pH, CaP-ppt content (Fig. 6a), and cell volume (Fig. 6b) of the culture supernatant of CJ cells (corresponding to the extracellular fluid on the lumen side) were measured at day-8 after varying the number of cells seeded on day-0. The higher the sown cell concentration, the lower the pH of the culture supernatant at day-8, and conversely, the higher the CaP-ppt content, with the pH at 2 × 10^4^ cells/200 μl/well seeding being almost 7.4 (Fig. 6a).

**Figure 6:**
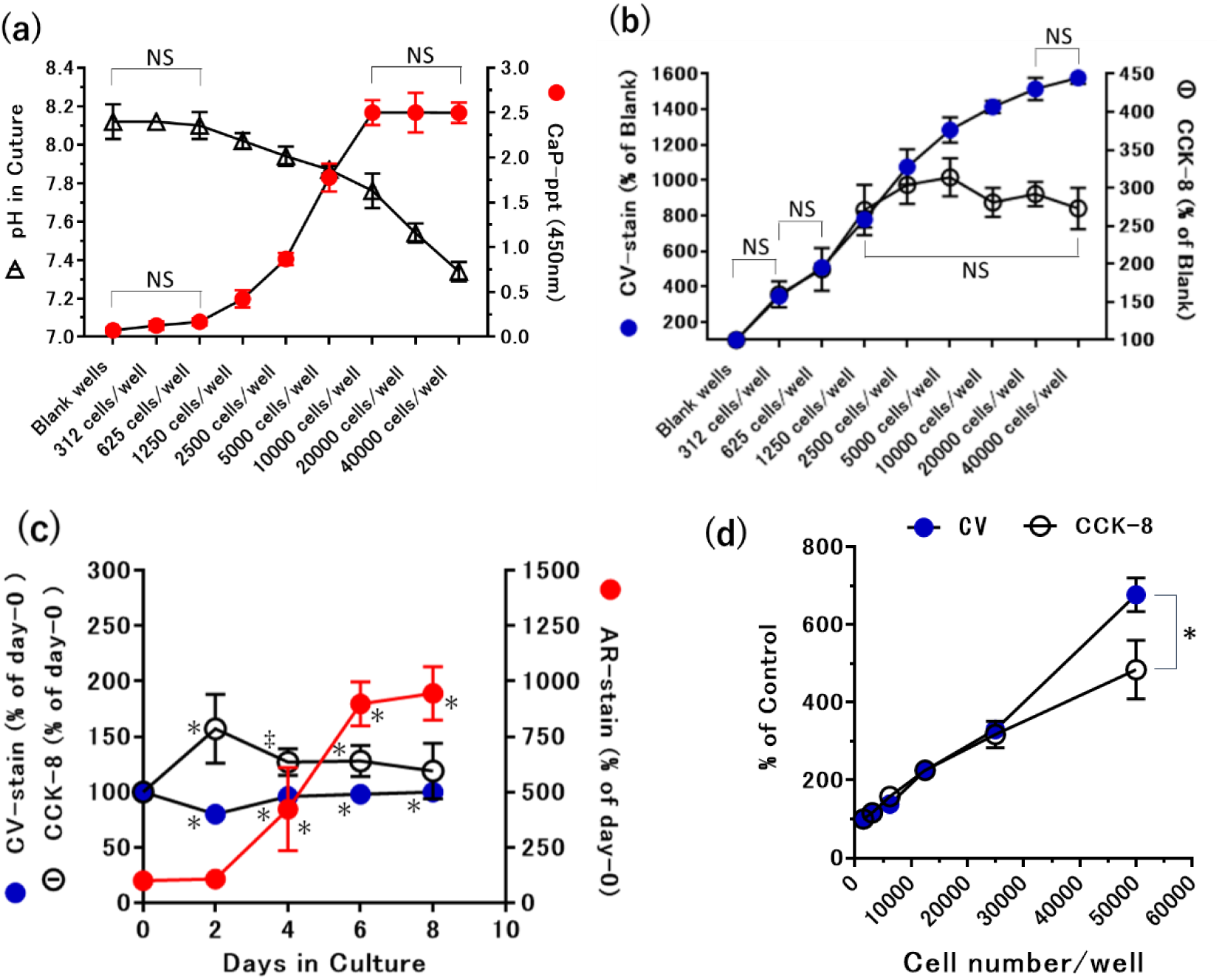
Active calcification of CJ cells in the physiological pH range. (**a**) and (**b**) On day-0, CJ cell dispersions at the indicated concentrations (n=8 per cell concentration) in 2-fold stepwise dilutions were dispensed at 200 μl/well and culture was initiated. On day-8, 20 μl/well of CCK-8 reagent was added first, and after 1 hour reaction in a CO2 incubator, OD 450 nm measurement was performed. The plate was returned to the CO2 incubator, and 2 hours later, pH measurement of the culture supernatant was quickly performed in 4 wells per cell concentration (n=4). During the pH measurement process, a film was attached to the top surface of other sample wells. Statistical analysis was performed by 2-way ANOVA with Tukey’s multiple-comparison test. The figure emphasizes and highlights the relationship with no significant difference between groups, while there is a significant difference of *p*<0.05 between groups without NS notation. (**c**) CJ cells were sown at 5 x 10^4^ cells/200 μl/well on day-0, and measurements were performed every 2 days: CCK-8 activity measurement, AR staining, and CV staining. The measurement on day-0 was performed 6 hours after cell adhesion and extension were confirmed. Statistical analysis was performed by 1-way ANOVA with Dunnett’s multiple-comparison test (n=8 for each measurement day) for day-0 (control). ‡*p*<0.005, \**p*<0.0001.(**d**) A 2-fold stepwise dilution of CJ cell dispersion with a maximum concentration of 5 x 10^4^ cells/200 μl/well was dispensed on day-0. After 6 hours, adhesion and extension of CJ cells to the plate was confirmed by specular inspection, additional 20 μl/well of CCK-8 reagent was added and After 1 hour of reaction in a CO2 incubator, the absorbance at OD450 nm was measured, and the CV staining was quantified. Statistical analysis was performed by 1-way ANOVA with Sidak’s multiple-comparison test on the corresponding number of cells. \**p*<0.0001.

Again, no signs of cell death were observed in this experiment, both measured with the CCK-8 kit and in the quantification of CV staining. However, above a certain amount of cell concentration, a difference appeared between these two assays, with the CCK-8 kit’s activity value not increasing above 1250 cells/well, whereas CV staining increased further with cell concentration (Fig. 6b). This difference between the two assays was expected to be a difference in detection of the incremental extracellular matrix produced by CJ cells. Therefore, we sowed high concentrations of CJ cells (5 x 10^4^ cells/well) on day-0 and examined calcification, CCK-8 activity, and CV staining intensity as a percentage change over time from day-0 (Fig. 6c). The expectation was slightly different. The expectation was that the temporal follow-up would show differences in CCK-8 activity and CV staining intensity like those in Fig. 6b, but there was neither a pattern of CV staining intensity exceeding CCK-8 activity nor an increasing trend. On the contrary, there was even an increase in CCK8 activity and a decrease in CV staining intensity (*p*<0.0001) on day-2 (Fig. 6c). The reason for these deviations from expectations, especially the failure to reproduce the increase in CV values, was presumably because the "6 hours" after seeding, when cell adhesion and elongation were complete, was the time point of day-0. In other words, something must have happened during this 6-hour period. Therefore, we compared CCK-8 activity and CV staining intensity after 6 hours of culture at different cell concentrations (Fig. 6d). As expected, at low cell concentrations before reaching confluency, both measurements were almost identical, but at 50,000 cells/well, CV staining intensity increased significantly (*p*<0.0001), diverging from the CCK-8 value.

### 3 – 6 : High-throughput screening (HTS) with CJ cells

Unlike rat VSMCs, the formation of CaP-ppt by CJ cells was not accompanied by cell death, and the medium pH 7.4 met the desirable pH conditions for calcification experiments as proposed by Hortells *et al.* [26]. Therefore, we performed the validation test as an HTS system for anti-calcifying substances by CJ cells. Fig. 7a is a photograph of the plate at day 8 after culture. Fig. 7b shows the pH value of the culture supernatant, which was 7.48±0.08 during CJ cell calcification compared to the SM row pH of 8.07±0.04 (n=6), an obvious significant difference (*p*<0.005).

**Figure 7:**
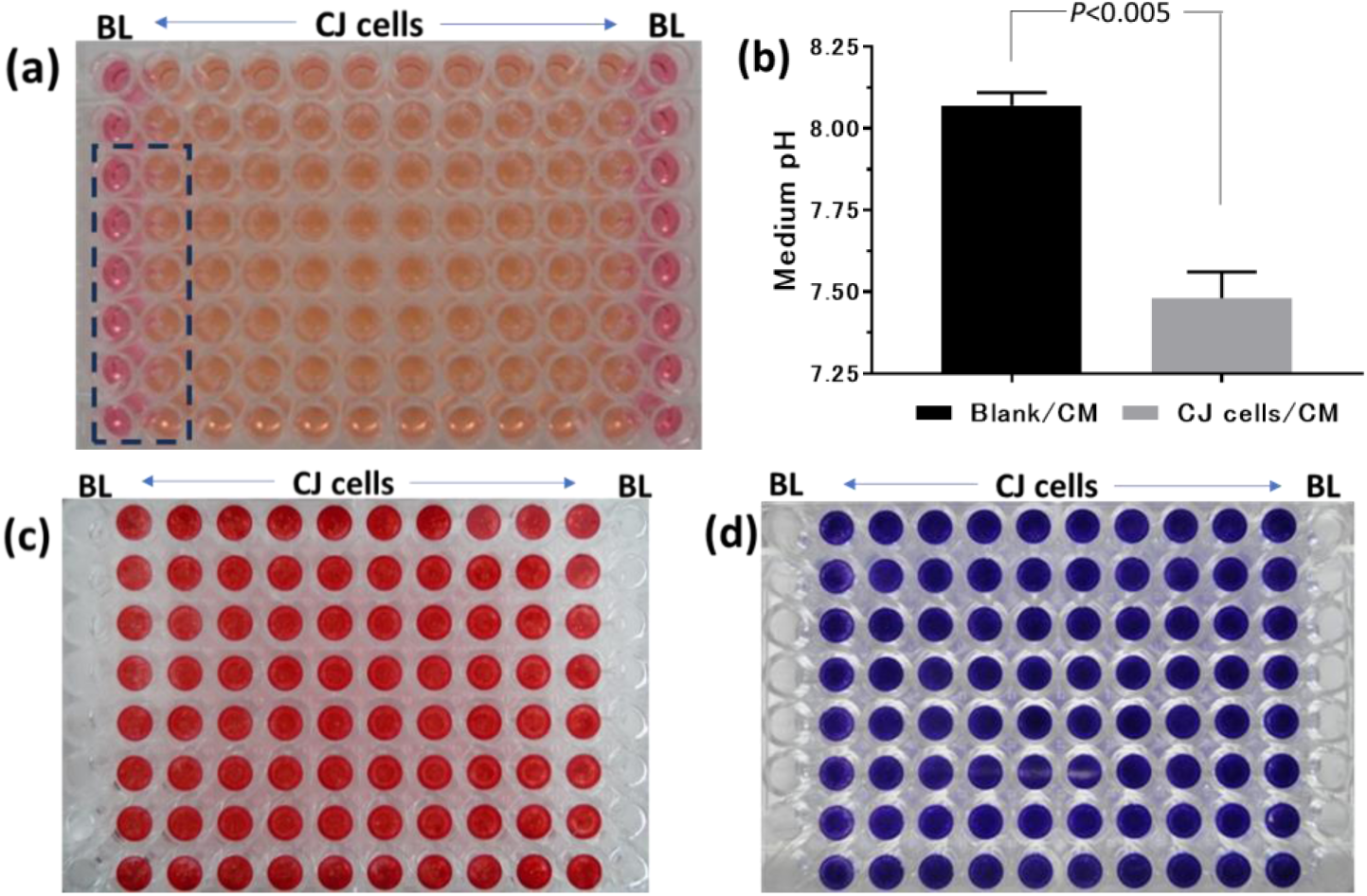
CJ cells show strong CaP-ppt formation ability in physiological Ca and phosphate concentrations and in the pH range and can be used as a screening system for calcification therapeutics. The validation assay for screening is described in the Methods section. (a) Phenol red coloration in the culture medium in the wells on day 10 of incubation. The remaining 80 wells containing CJ cells show an orange-neutral color, whereas the CJ-free wells (blank) at the left and right ends show a purple-alkaline color. The medium was not changed at all. (b) Results of pH measurement in the two dotted rows on the left side of (a) (n=6 each). (c) is a photo of a dried plate after AR staining, taken from the bottom and reversed left to right from (a). (d) Photograph of CV-dyed dried plate on the same plate. BL: Blank column.

Fig. 7c is an AR stained (CaP-ppt) image of the same plate, and Fig. 7d is a CV stained (cells) image performed on the same plate. In both stains, the blanks at both ends were completely unstained. Table 2 summarizes the parameters of the validation experiments, which were performed twice with the determination days as day-8, day-10, and day-12. All parameters were at satisfactory levels and the evaluation system was robust.

**Table 2:** Robust anti-calcification screening parameter values for CJ cells. CJ cells were cultured under the conditions described in Fig. 7, with no medium change during the period. AR and CV stains were performed on days 8, 10, and 12 in three independent 96-well plates; CaP-ppt: calcium phosphate precipitation; Co-Va: coefficient of variation; S/B: signal-to-background ratio; S/N: signal-to-noise ratio. Z’-factor: a measure of the degree of optimization of the assay system. *^)^ In the CV stain after 8 days, as shown in Fig. 6, not only nucleic acids of living cells but also ECM (extracellular matrix) consisting of a considerable amount of proteins, lipids, and other macromolecules are co-stained, but the evaluation of the anti-calcification potential of a compound is concerned with the presence and degree of cytotoxicity. Therefore, the term "cells" is used here to include the organic materials.

| Trial | Parameter | Acceptability | CaP-ppt (OD450) on ; |  |  | Cells*(OD590) on ; |  |  |
| --- | --- | --- | --- | --- | --- | --- | --- | --- |
|  |  |  | day-8 | day-10 | day12 | day-8 | day-10 | day-12 |
| First | Co-Va | <10% | 8% | 7% | 8% | 4% | 4% | 4% |
|  | S/B | >3 | 49 | 51 | 49 | 16 | 20 | 19 |
|  | S/N |  | 281 | 487 | 421 | 124 | 350 | 167 |
|  | Z'-factor | >0.5 | 0.7 | 0.8 | 0.8 | 0.9 | 0.9 | 0.9 |
| Second | Co-Va | <10% | 7% | 8% | 8% | 4% | 4% | 3% |
|  | S/B | >3 | 46 | 53 | 50 | 20 | 21 | 22 |
|  | S/N |  | 308 | 171 | 509 | 242 | 210 | 345 |
|  | Z'-factor | >0.5 | 0.8 | 0.7 | 0.8 | 0.9 | 0.9 | 0.9 |

### 3 – 7 : Anti-CaP-ppt action of bisphosphonates

Etidronate is a first-generation bisphosphonate that has been reported to be effective in preventing [37] and curing [38] arterial calcification in human clinical practice.

Alendronate, a nitrogen-containing bisphosphonate, has also been reported to be effective in an *in vitro* renal stone model using MDCK cells [39]. These bisphosphonates were compared and evaluated in the CJ cell HTS system, and the results are shown in Fig. 8. The IC_50_ for the anti-CaP effect of alendronate is 2.6 μM, but the IC_50_ for the cytotoxic effect is also 10.9 μM, and the IC_50_ ratio of the two is close at 4.2-fold (Fig. 8b). Rather, etidronate, a first-generation bisphosphonate, had a weaker IC_50_ value of 39.2 μM for its anti-CaP effect than alendronate, but showed almost no cytotoxicity and a high safety profile with an IC_50_ value ratio of approximately 100-fold between the two.

**Figure 8:**
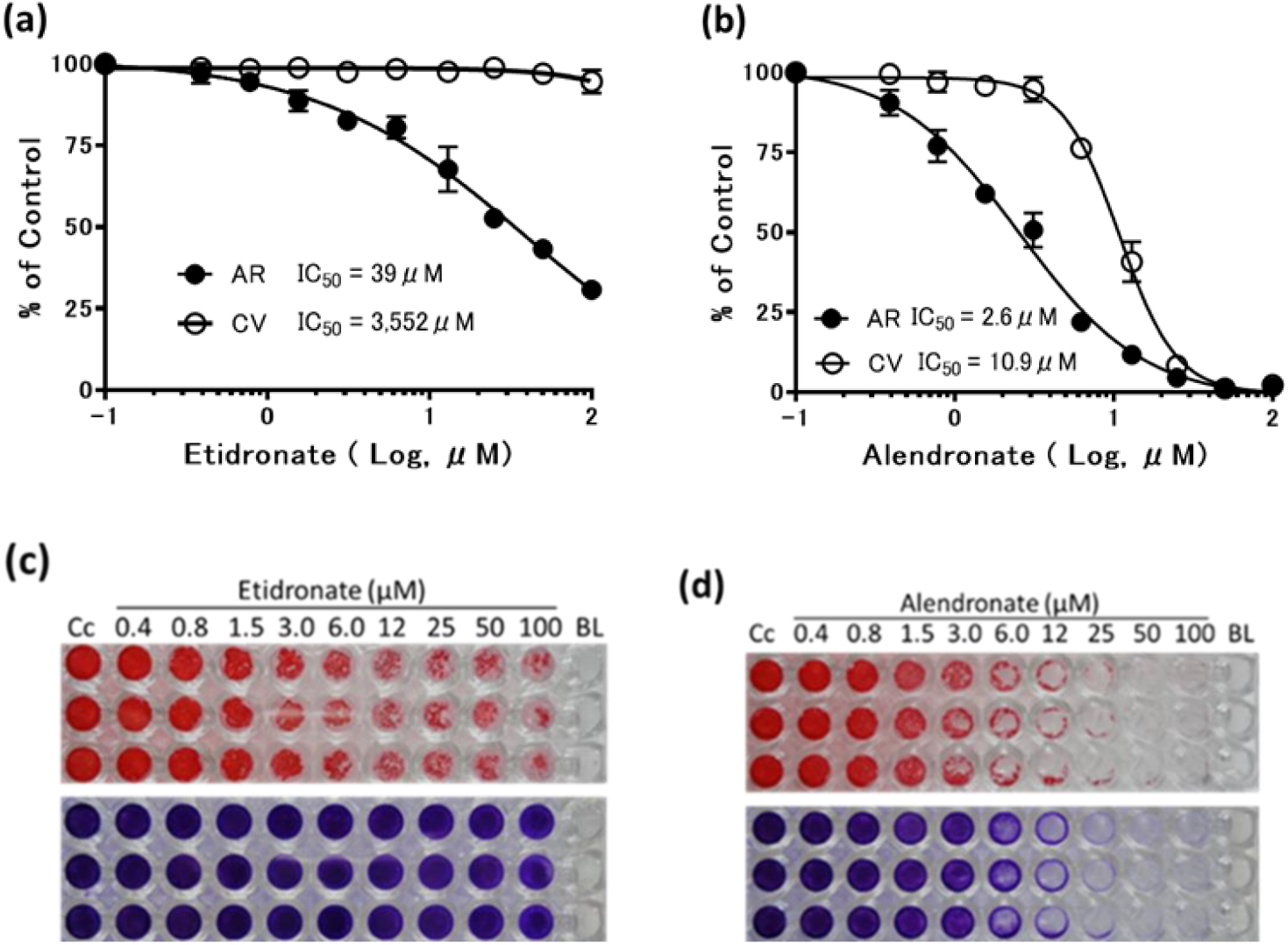
Anti-calcification effect of bisphosphonates in the CJ cell screening system. The method was performed as described in the Methods section. (**a**) Dose-response effect of etidronate (top panel) and (**c**) stained plate photograph (bottom panel). (**b**) Dose-response effect of alendronate (upper panel) and (**d**) stained plate photograph (lower panel); AR: Alizarin Red S staining, CV: Crystal Violet staining, Cc: cell control, BL: blank.

### 3 – 8 : Anti-CaP-ppt action of all-trans retinoic acid (ATRA)

Next, we evaluated ATRA, which has been reported to be effective in calcification of heart valves and arteries [40], with CJ cell HTS. Fig. 9a and 9b are stained images of AR- and CV-stain, respectively. Fig. 9c shows the measurement of medium pH on the day of determination, and the anti-CaP-ppt effect of ATRA occurred at physiological pH. The very strong anti-calcification effect of the IC_50_ value of 0.14 μM for AR staining in Fig. 9d was satisfactory from the plate staining image. However, the IC_50_ value of 742 μM for CV staining was disconcerting based on the staining diagram and dose-dependence curve, and the results of Tukey’s multiple comparison test using the CV staining value as 100% of the OD 590 nm value without ATRA. Indeed, in quantification, there was a significant difference in CV staining intensity between ATRA concentrations of 0.312 μM and 0.624 μM (*p*<0.05), as shown in Fig. 9e. However, there was no difference between subgroups below 0.312 μM or above 0.624 μM. This strange discontinuity was more prominently replicated in the retest (Fig. 9f) (*p*<0.0001).

**Figure-9 :**
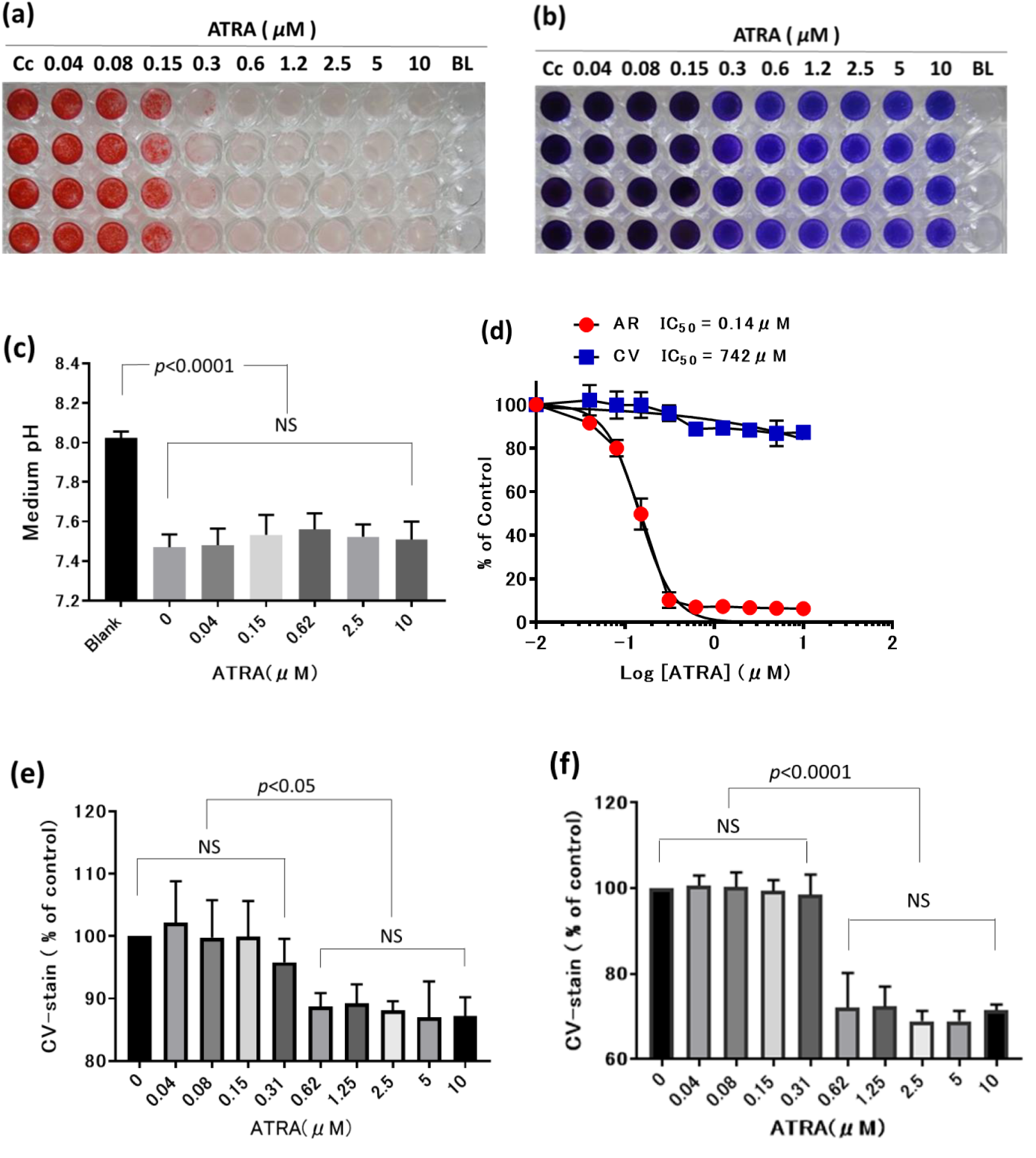
Anti-calcifying effect of all-trans retinoic acid (ATRA). (**a**) AR staining image of the anti-calcification effect of ATRA, (**b**) Color change in CV stainability in the same test, (**c**) pH value of the culture medium on the day of determination, (**d**) IC50 of AR staining intensity and slight decay of CV staining intensity by ATRA, (**e**) Discontinuous decay response of ATRA on CV staining intensity after one 5% formic acid extraction in the usual way. Tukey’s multiple comparison test results between groups at each concentration (n=4). (f) Tukey’s multiple comparison test results between groups at each concentration (n=4) after 3 extractions with 5%-formic acid. Staircase-like CV staining intensity change is more obvious between ATRA concentration 0.31μM and 0.64μM. BL: Blank column. Cc: Cell control

## 4 : Discussion

Unlike our original purpose, this study is a characterization of the unique properties of an accidentally collected canine renal adenocarcinoma cell line (CJ line) with spontaneous and potent calcification capacity. After conducting a literature review, we concluded that CJ cells have the potential to be a new research material that can solve the problems of conventional research material regarding RSD or CVC, which are representative of ECD. The problems are the weak calcification signal in conventional *in vitro* cellular model systems and, especially in CVC diseases, doubts about the validity of the *in vitro* hyperphosphatemic culture system as a model based on hyperphosphatemia in CKD and ESRD patients [26, 29].

ECD include aging, diabetes, kidney stones, Fahr’s disease, age-related macular degeneration, breast cancer, pancreatic cancer, prostate cancer, ovarian cancer, thyroid cancer, systemic sclerosis, calcific tendonitis, and Fahr’s disease [2]. In these diseases without hyperphosphatemia, the mechanism of early calcification in the patient’s body is largely unknown. Thus, CJ cells, which exhibit spontaneous and potent calcification in cultures prepared to mimic healthy human blood or body fluids, can be considered an *in vitro* model for many ECD.

As already noted, there are few reports of spontaneously calcifying cell lines, with only the CVC strain [30, 31] and MDCK strain [32, 33], and even in studies using the CVC line, calcification was subsequently promoted by adding *β*-GP at concentrations of 2.0∼5.0 mM [41, 42]. In both cases, there are few issues that can be compared or discussed with the results of the CJ cell experiments, other than the differences in the intensity of the calcification signal. Therefore, the results of this study are also a comparative discussion of many high-concentration phosphate-induced experimental cases and a few spontaneous calcification cases. First, we will again summarize the results of this experiment and discuss each issue separately.

CJ cells had epithelial-like morphology (Fig. 1b,c), doubling time was about 24 h (Fig. 1e), and the average chromosome number approximated the haploid number of 78 for normal canine cells [43] (Fig. 1f). Spontaneous calcification began after reaching confluence and grew as CaP particles of unequal size, aggregates, and fusions (Fig. 1d), as well as in plate, linear, and other forms (Fig. 2); CaP-ppt formation was FBS concentration dependent (Fig. 4), with a Ca/P ratio of 1.35 and a constitutive ratio of ACP2 (Fig. 5c); Ca and Pi in the CaP-ppt were derived from Ca^2+^ and PO_4_^3-^ ions in the supernatant culture medium (Fig. 5a,b), suggesting that some sort of feed-back regulation mechanism was triggered by the CJ cells as both ions in the supernatant decreased, and Ca^2+^ and PO_4_^3-^ in the supernatant were not depleted (Fig. 5a,b). CJ cell survival during CaP-ppt formation was also demonstrated by the commercial CCK-8 kit (Fig. 6b,c), and CaP-ppt formation proceeded at physiological pH (neutral) (Fig. 6a, Fig. 7). Anti-CaP-ppt compound screening could be evaluated in combination with cytotoxicity testing on the same plate (Fig. 7c,d), and a low-cost and reliable HTS system was established (Table 2). Two bisphosphonates (Fig. 8a,b) and ATRA (Fig. 9) were evaluated in this CJ cell HTS system, and results were consistent with human clinical and animal studies of these compounds, as discussed below.

First, the effect of FBS on calcification is discussed in the CJ and cardiovascular cell systems. In conclusion, the requirement for FBS is not constant: CJ cells showed a distinct FBS requirement in CaP-ppt formation (Figure 4). Similar FBS-dependent CaP-ppt formation has been reported in an *ex vivo* organ culture system with 3.8 mM *β*-GP-added rat aortic rings [44, 45]. On the contrary, it has been reported that FBS suppresses calcification in planar culture systems of rat VSMCs [26] or human VSMCs [44] with *β*-GP added. However, the reason for the FBS-induced calcification of CJ cells is still unclear.

Next, the presence of cell death during calcification is discussed. In the calcification of arterial SMC, the involvement of cytotoxicity due to early-forming CaP particles associated with alkalinization of the culture medium was demonstrated [26]. It is not surprising that even in calcification of CJ cells of renal origin, cell death could theoretically be caused by early-formed CaP-microparticles. This is because calciprotein monomer (CPM), an early product of CaP-ppt, and its aggregate, primary calciprotein particle (CPP), are cytotoxic to human renal proximal tubular cells (HK-2 strain) [46]. It has even been postulated that these two ACPs may be the main body of uremia [47]. Given the large amount of CaP-ppt produced by CJ cells, a large amount of cytotoxic ACP should have been produced even in the early phase, and CJ cells should have died, but the reality was quite different. The CCK-8 kit, which evaluates viable cell numbers via mitochondrial enzyme activity, showed no signs of cell death (Fig. 6b,c), and the trypan blue dye exclusion test also confirmed survival (Fig. 1e). Furthermore, medium acidification due to active cell metabolism during culture also supported the presence of viable cells (Fig. 6a). The reason for this difference in performance is still unknown.

The solution pH problem during calcification is then discussed. In general, there is a traditional view of the EC phenomenon as a cellular degenerative, passive process and a newer view as an active, proactive process mediated by cellular metabolism [45]. The most widely used experimental system for CVC is the combination of primary cultures of arterial smooth muscle or valve stromal cells with high concentrations of phosphoric acid [19]. The above-mentioned cell death by high concentrations of phosphate inevitably leads to the loss of cellular metabolic systems, and CaP-ppt formation was explained as a physicochemical reaction associated with the alkalinization of the medium pH by high concentrations (44 mM) of sodium bicarbonate in DMEM culture medium [26]. In other words, this general-purpose experimental system has observed a passive process rather than an active one. In contrast, CJ cell calcification is not a passive reaction, as there is neither cell death nor media alkalinization. On the contrary, CJ cell calcification is a good example of a truly active process that maintains a physiological pH around 7.4 via acid production, a result of high metabolic capacity (Fig. 6a,b,d). The reasons for active formation of calcium deposition at a high energy cost are interesting and need to be pursued separately and seriously. For example, the suggestion of the existence of a kind of feed-back regulation mechanism that maintains cell survival without depleting Ca^2+^ and PO_4_^3-^ ions in the culture supernatant (Fig. 5a,b) is also very interesting and may provide clues to the clarification of active processes.

Of course, it is possible that *in vivo* calcification is controlled by local alkalinization of the body environment [26, 29]. Already in 2009, De Solis *et al.* conducted experiments in CKD patients on dialysis with the hypothesis that alkaline load is involved in arterial calcification. Their results showed that simply increasing the pH from 7.42 to 7.53 in a *β*-GP-added VSMC culture system resulted in a 2.5-fold increase in CaP-ppt formation, and sodium bicarbonate administration to 5/6 nephrectomized mice resulted in a threefold increase in aortic calcification index [48]. Even in bone metabolism, a physiologically calcified tissue, the addition of high concentrations of phosphoric acid (10 mM *β*-GP) caused calcification of mouse osteoblasts MC3T3-E1 under alkaline (pH 7.8) but not under physiological pH (pH 7.4) conditions [49]. In addition, corals have been observed to generate a higher pH environment in their bodies than in the surrounding seawater to maintain calcification and counteract the threat of dissolution of CaCO_3_ shells during seawater acidification caused by increased atmospheric CO_2_ gas [50, 51].

On the other hand, we must assume the existence of a hidden mechanism of calcification in the body, which is non-hyperphosphatemic, non-cellular toxic, and non-alkalinizing. This silent calcification mechanism may be present in the early stages of CKD, ESRD, and even in healthy individuals as described below, as well as in various EC diseases without hyperphosphatemia. Ca-ppt formation in CJ cells is highest near neutral, the physiological pH, and Ca-ppt formation is lowest on the alkaline side [26, 29], where the concentration of PO_4_^3-^, the only phosphate ion species that makes up ACP and HAP, is highest (Fig. 6a, Fig. 7a,b). Above all, the calcification of CJ cells is unique in that they emit a strong calcification signal at physiological phosphate concentrations (0.9 mM), which is unprecedented. Clearly, CJ cells have the potential to be an interesting research material for elucidating the silent calcification mechanism in both basic and applied studies of ECD.

Furthermore, the calcification kinetics of CJ cells are discussed. The total amount of Ca^2+^ and PO_4_^3-^ allocated to the culture supernatant and CaP-ppt was within the theoretical range after 6 days of culture, but 20%∼30% less than the theoretical value early in the culture (day 2, day 4) (Fig. 5a,b). A simple reason for this may be that after supernatant collection, the wells were washed twice with Hepes buffered saline before EtOH fixation, and these washing operations may have resulted in the loss of the unfixed CaP-ppt. In other words, at this early stage of culture initiation, poor adhesion of ACP1 and other Ca particles generated early in the process to the cell layer is suspected. Indeed, the fact that ECM (Fig. 9e,f) production, which is suggested to be involved in the adhesion and uptake of CaP particles, is enhanced in the later stages of culture (Fig. 6b,d) also supports this presumption. This interesting observation of materials balance was first obtained with the use of CJ cells. In other words, osteoblasts [12], vascular smooth muscle cells [19], and cardiac valve stromal cells [20] all require medium changes every 2 to 3 days during culture, making it difficult to obtain accurate material balance data in the open systems. On the other hand, CJ cells can be cultured for a long period of time without medium change in the closed system, which enables accurate tracking of material balance. The advantage of the CJ cell system, which does not require labor-intensive and time-consuming medium exchange, is very great when drug screening, as described below, is considered.

In general, calcification of the heart and blood vessels progresses over decades [6, 52]. Intimal thickening, a precursor lesion to atherosclerosis, forms in humans as early as 2 years of age, and in almost everyone by the age of 10. Intimal thickening is an adaptive response to the forces (blood pressure) on the arterial wall and is considered an inevitable change with age [53]. In case of RSD the average formation time of kidney stones was 5 years, with the slowest taking as long as 23 years [54]. In other words, even in a seemingly normal body, the initial lesion (etiology) is already present. Ironically, episodes of kidney stone formation are said to be a "red flag," a warning sign of arteriosclerosis that has been quietly progressing for many years [55]. Interestingly, in the detection of this very early lesion, there are older studies that have historically been accumulated in the RSD area than in the CVC area.

There have been a series of pioneering reports in which CaP particles were normally detected in the kidneys of healthy individuals [56, 57, 58]. Bruwer followed up on these reports and proposed that CaP particles in the kidneys of healthy individuals, including his own x-ray data, could lead to Randall’s plaques [59] . Brewer’s “Anderson-Carr-Randall progression theory” has been supported by recent advanced imaging equipment such as high-resolution μCT, electron microscopy, and correlation microscopy [60, 61, 62, 63]. In fact, early Randall’s plaques were detected in 73% of healthy renal papillae with the presence of ACP at a high rate of 80% [64]. ACP can be medically treated for early Randall’s plaques.

Furthermore, the scenario in which kidney stones grow with increasing size by the pathway of Ca^2+^, PO_4_^3-^ ions ⇒ CaP plate ⇒ CaP cluster ⇒ calcium nanoparticle (CNP) ⇒ ACP ⇒ Randall’s plaque is also almost established. Therefore, it was natural to emphasize that the target of drug therapy should be upstream of the Randall’s plaque, before ACP formation [62, 65, 66]. Scanning electron microscopy and energy-dispersive X-ray spectroscopy revealed that CaP particles ranging in size from 0.1 μm to 5 μm (mean 0.82 μm ± 0.67 μm, n=763) are commonly present in healthy heart valves [67, 68]. Van Engeland *et al.* argued that CaP particles, which were also found in heart valves of healthy individuals, were not innocent bystanders but induced pathological changes in the vascular intima and stromal cells and should be the subject of therapeutic studies [68]. *In vitro* CJ cell spontaneous calcification system contains CaP particles (Fig. 2) of the size of those in healthy individuals. Of course, the closed culture system of CJ cells must have undergone the earliest CaP-ppt particulate formation step between Ca^2+^ and PO_4_^3-^ ions to ACP2. Therefore, CaP-ppt particles, which are collected from patients and healthy subjects as research material [67, 68], could be prepared in large quantities from CJ cells. Thus, there is a linkage and consistency between *in vivo* (living organisms) and *in vitro* (cultured cells) based on the "common axis of spontaneous calcification.

Next, we discuss the quality of CaP-ppt formed by CJ cells in an in vitro culture system. Until now, the gold standard for determining the quality of CaP-ppt, i.e., Ca/P ratio, has been X-ray diffraction and Fourier transform infrared spectroscopy. However, data comparable to those results can also be obtained with commercially available Ca and Pi measurement kits[25].The Ca/P calculation results obtained from commercial kit measurements showed that the quality of the CaP-ppt formed by CJ cells was ACP2 (Fig. 5c); ACP2 is also a major advantage in terms of conducting large-scale screening and basic research. In medical intervention, ACP2 is an ideal target for prophylactic and therapeutic drug discovery before the point of no-return [5, 6, 29], since the CaP-ppt formed is a form that can revert to soluble Ca and P [29]. Furthermore, the fact that the Ca/P ratio remains constant from 6 to 14 days of incubation provides stability and reliability in the screening of anti-CaP compounds. Indeed, validation with CJ cells showed that all parameters were acceptable (Table 2). Higher-order screening, e.g., in conjunction with animal studies, is also important: Hortells *et al*. performed *in vivo* arterial calcification induction experiments in 5/6 nephrectomized rats plus high-phosphate diet feeding (a model of stage 3-4 CKD), which are more reliable than *in vitro*. Particle size and its Ca/P ratio in rat VSMC were measured using a field emission scanning electron microscope equipped with an energetic dispersive spectroscopy system. The results showed that the CaP particle size was 0.1∼0.4 μm and the Ca/P ratio was 1.35 in ACP2 [69].That is, even in the *in vitro* CJ cell system, CaP particles less than 1 μm in size are generated (Fig. 1d, Fig. 2), and the Ca/P ratio is 1.35 ACP2, which is consistent with the *in vivo* results of Hortells *et al*. Furthermore, from the basic research aspect, what kind of stimulus is needed to progress from ACP to HAP, the point of no-return? This research approach and results will also be a target [29] for the search for therapeutics (therapies that slow the progression to HAP or re-dissolve HAP) as mentioned by Lanzer.

Whether the therapeutic target is ACP or HAP is an important point [6, 29] in the development of anti-CaP drugs. That is, up to ACP, CaP-ppt can return to the soluble state of Ca and P, but when it progresses to HAP, it becomes irreversible. Fresh CaP-ppt formation appears as a low-density sheet called ACP1, with a Ca/P ratio of 1, followed by spherulite called ACP2, with a Ca/P ratio of 1.35 as maturation proceeds, and finally HAP with a Ca/P ratio of 1.67 as maturation proceeds [26, 29]. Therefore, the transition point from ACP to HAP is a "point of no-return," requiring a switch in perspective from prevention up to ACP to treatment when HAP is reached [6, 29]. However, as mentioned above, there are currently no approved drugs for either prevention or treatment points.

The use of CJ cells is useful as a drug screening system, which is an urgent issue. The system is an active reaction with live cells that calcifies at physiological pH around 7.4 (Fig. 7) and shows highly reliable Z’-factor values (Table 2). Furthermore, the system does not require medium exchange, and both calcification and cytotoxicity can be evaluated in the same plate [4, 6].

First, we were interested in bisphosphonates and ATRA and tested them in the CJ cell HTS system (Fig. 8 and 9). While both cardiovascular disease and kidney stones progress over many years, generalized arterial calcification in infancy (GACI) is a short-lived disease that is hereditary and results from a systemic defect in nucleotide pyrophosphatase (NPP1) activity (E.C. 3.6.1.9). The systemic deficiency of vascular occlusion (E.C. 3.6.1.9) results in a devastating ECD in which most affected children die in early infancy from congestive heart failure due to myocardial infarction or hypertension. A study of the first-generation bisphosphonate etidronate against GACI resulted in a reduction in arterial calcification and a doubling of survival [38]. Inhibition of arterial calcification and retinal neovascularization was reported as a result of etidronate treatment of EC of elastic fiber pseudoxanthoma [37]. On the other hand, a report showed that third-generation alendronate, which was expected to have a stronger effect, did not change aortic calcification in 5/6 nephrectomized rats and CKD dialysis patients [70], while etidronate suppressed aortic calcification [71], suggesting that the nitrogen content of bisphosphonates makes a difference. In addition, there are negative reports of atrial fibrillation, angina pectoris, coronary atherosclerosis, and arrhythmogenesis with alendronate in humans[72], and rupture of atherosclerotic plaques in Apo-E-/- mice[73]. The IC_50_ value of etidronate is higher, but the anti-CaP-ppt effect is sufficient with little cytotoxicity (Fig. 8a,c). On the other hand, although the anti-CaP action of alendronate was stronger than that of etidronate, it also exhibited a strong level of cytotoxicity (Fig. 8b,d). Thus, the anti-CaP-ppt formation inhibitory effect of alendronate is an apparent cytotoxic effect, indirectly supporting active CaP-ppt formation in CJ cells and also explaining its inferior anti-calcification effect compared to etidronate in vivo. Incidentally, risedronate also showed similar cytotoxicity to that of alendronate in the CJ cell evaluation system (data not shown).

ATRA has been reported to inhibit calcification of human coronary artery SMC cultured in DMEM with 10 mM *β*-GP [40] and to inhibit intra chondral heterotopic ossification in mice [74] . In case of the anti-CaP-ppt effect in our CJ cell HTS, ATRA exhibited a strong anti-CaP-ppt effect with an IC_50_ value of 0.12 μM. Interestingly, as shown in Fig. 9b, the CV staining intensity was not continuous, with a significant difference (*p*<0.05) between the low and high ATRA concentrations, indicating a threshold between 0.31 μM and 0,62 μM. This discontinuous and dramatic color change was further accentuated (*p*<0.0001) in the retest using a stronger formic acid washing. This suggests that extracellular matrix (ECM) production is involved in the spontaneous calcification phenomenon of CJ cells, and that the anti-CaP-ppt effect of ATRA is due to inhibition of ECM production. CV dyes stain not only nucleic acids but also proteins, lipids, and all other macromolecules (Nilles), but the actual ECM involved in CaP-ppt formation in these CJ cells is unknown. What is most interesting is the extreme divergence of staining at a certain threshold, as if the ATRA-treated CJ cells underwent a simultaneous phase transition to cells with completely different properties. Hutchson & Goettsch describe that "therapies developed for CKD patients may be applicable to other patients at risk for developing EC" [6] . Of course, conversely, even kidney-derived CJ cells could be used for prophylactic and therapeutic drugs for CVC diseases.

In conclusion, the conventional model of EC with high concentrations of phosphate is questionable, but it is still a viable model if there is local alkalinization of the lesion. However, it is unlikely to be an in vitro model for other ECD that do not involve hyperphosphatemia.

On the other hand, CJ cells can serve as an *in vitro* model for most ECD. The Ca/P ratio of CaP-ppt produced spontaneously by CJ cells is 1.35 of ACP, and its practical application is as a drug discovery system for the early stage (reversible stage) of ECD, where patients are in dire straits and waiting for an effective drug. In addition, the system has the potential to meet the expectations of patients who are still waiting for an effective drug in dire circumstances. Furthermore, why do organisms that minimize waste and conserve energy pay a high energy cost to form ECCJ cells are expected to be a useful research material that can be developed and expanded into various research themes other than the field of EC due to their various unique characteristics.

## Acknowledgements

We thank the following members for providing tumor material. The Fund member doctors (K. Takashima, T. Yamane, J. Ogasawara, Y. Mizutani, Y. Kaneko-Wada, M. Kawasaki, S. Tanno, M. Sakamoto), and the following doctors: T.Tezuka (Tezuka Animal Hospital), K. Koide (Ikasa Animal Medical Center Koide Animal Hospital), K. Kato (Kato Animal Hospital), M. Uno (Central City Animal Hospital), M. Takenaka (Takenaka Animal Hospital), M. Sato (Sato Veterinary Clinic), and K. Noro (Noro Animal Hospital). We thank Dr. J. Itoh (Tokai University) for the provision of MDCK cells, Dr. N. Itoh (Tottori University) for convenience in obtaining experimental equipment, and Mrs. M. Yoshida for experimental support.

## Author Contribution

N.I. and Y.Y. designed the study. N.I. performed *in vitro* experiments and analyzed data. Y.Y. was involved in the manuscript revision and data analysis.

## Abbreviations and Acronyms (alphabetical order)

ACP: amorphous calcium phosphate
*β*-GP: beta-glycerophosphate
CaP-ppt: calcium phosphate precipitate
CKD: chronic kidney disease
CVC: cardiovascular calcification
EC: ectopic calcification
ECD: ectopic calcification disease
ESRD: end stage renal disease
HAP: hydroxyapatite
HTS: high throughput screening
RSD: renal stone disease
S: supersaturation
SM: Dulbecco’s modified Eagle’s minimum essential medium containing 10% FBS, 100 u/ml penicillin, 100 μg/ml streptomycin, and 4.5 g/L glucose
VSMC: vascular smooth muscle cells

